# A chromosome-level genome assembly and annotation of the humpback grouper *Cromileptes altivelas*

**DOI:** 10.1101/2020.06.22.164277

**Authors:** Yun Sun, Dongdong Zhang, Jianzhi Shi, Guisen Chen, Ying Wu, Yang Shen, Zhenjie Cao, Linlin Zhang, Yongcan Zhou

## Abstract

*Cromileptes altivelas* that belongs to Serranidae in the order Perciformes, is widely distributed throughout the tropical waters of the Indo-West Pacific regions. Due to their excellent food quality and abundant nutrients, it has become a popular marine food fish with high market values. Here, we reported a chromosome-level genome assembly and annotation of the humpback grouper genome using more than 103X PacBio long-reads and high-throughput chromosome conformation capture (Hi-C) technologies. The N50 contig length of the assembly is as large as 4.14 Mbp, the final assembly is 1.07 Gb with N50 of scaffold 44.78 Mb, and 99.24% of the scaffold sequences were anchored into 24 chromosomes. The high-quality genome assembly also showed high gene completeness with 27,067 protein coding genes and 3,710 ncRNAs. This high accurate genome assembly and annotation will not only provide an essential genome resource for *C. altivelas* breeding and restocking, but will also serve as a key resource for studying fish genomics and genetics.

## Data Description

### Background & Summary

The humpback grouper *Cromileptes altivelas* (order Perciformes, family Epinephelinae) inhabits the tropical waters of Indo-West Pacific oceans^1^. *C. altivelas* is increasing attracting attention as high-value human food for its delicious flavor and high nutritional value, and it also has great ornamental value due to its unique body shape and beautiful color^1-3^ (Fig.1).However, the wild population of *C. altivelis* is increasingly exploited. Meanwhile, *C. altivelis* farming is limited by its slow growth speed, low survive rate, and various pathogenic diseases^4-5^. Obtaining high-quality genomic sequences is the foundation of developing genomic selection to improve the performance of *C. altivelis*. The genome information is also critical to explore the genetic mechanisms of its unique traits, immune system and evolutionary adaptation. Recently, genome sequences of seven grouper fish species are available. Most of these fish species belong to the genus of *Epinephelus*. There are few genome sequences of grouper fish species from other genera. Humpback grouper is the only species of *Cromileptes* genus.

Here, combining a PacBio long-read sequencing and high-throughput chromosome conformation capture (Hi-C) technologies, we sequenced the humpback grouper *C. altivelas* genome with estimated size 1.07 Gb. The N50 scaffold size of final genome assembly reached 44.78Mb and 99.24% of the scaffold sequences were anchored into 24 chromosomes. Based on the high-quality assembly, we annotated the protein-coding genes and ncRNAs. The high-quality genome assembly and annotation will not only provide an essential genome resource for exploring the economic values of *C. altivelas* breeding and restocking, but will also serve as a key resource for studying fish genomics and genetics

## Methods

### Sample collection, library construction and sequencing

We sampled a single individual of female *C. altivelas* for genome sequencing from Hainan, China (Fig.1). The total genomic DNA was extracted from muscular tissue using SDS lysis and magnetic beads isolation method.

We applied a strategy combing four technologies for library construction and sequencing including PacBio Sequel System (for genome assembly), the Illumina Hiseq 4000 System (for genome survey), 10X Genomics link-reads (for scaffold construction), and Hi-C optical maps (for chromosome construction). First, two paired-end Illumina sequence libraries were constructed with an insert size of 350 bp, and sequencing was carried out on the Illumina HiSeq 4000 platform. A total of 79.18 Gb (coverage of 71.98 X) of Paired-End 150 bp reads were produced. Raw sequence data generated by the Illumina platform were filtered by the following criteria: filtered reads with adapters, filtered reads with N bases more than 10%, and filtered reads with low-quality bases ⩽5) more than 50%. Second, a total of 113.49 Gb of polymerase reads data were generated using PacBio Sequel platform, and a total of 106.3 Gb (coverage of 103 X) subreads were obtained after removing adaptors and filtered with the default parameters. The average and the N50 length of subreads reached 8.04 kb and 13.26 kb, respectively. Third, one 10X Genomics linked-read library was constructed and sequenced on Illumina HiSeq 4000 platform, which produced 129.1Gb (coverage of 117.4 X). Finally, an optical map was also constructed from Hi-C, of which 119.2 Gb (coverage of 108.4 X) data were generated. All sequence data are summarized in Table 1.

**Table 1.** Summary of sequencing data generated in this study.

| Pair-end libraries | Insert size | Total data (G) | Read length (bp) | Sequence coverage (X) | Application |
| --- | --- | --- | --- | --- | --- |
| Illumina reads | 350 | 79.18 | 150 | 71.98 | Genome survey |
| Pacbio reads | - | 113.49 | - | 103.17 | Genome assembly |
| 10X Genomics | - | 129.1 | - | 117.4 | Scaffold construction |
| Hi-C | 350 | 119.2 | 150 | 108.4 | Chromosome construction |
| RNA-seq |  | 38.67 | 150 | 35.2 | Genome annotation |

Note: The coverage was calculated using an estimated genome size of 1.07 Gb.

### Genome size estimation

The genome size of *C. altivelas* was first estimated using *k*-mer spectrum with Jellyfish^6^ (v2.1.3). The distribution of 17-kmer showed a major peak at 57 (Figure S1). Based on the total number of kmers (63,765,804,944) and corresponding to a kmer depth of 57, the *C. altivelis* genome size was estimated to be 1118.70 Mb using the formula: Genome size= kmer_Number / Peak_Depth. The modified genome size was 1104.81 Mb, the genome heterozygosity was 0.16%, and the repetition rate was 46.38%.

### *De novo* assembly of the *C. altivelis* genome

The contig assembly of the *C. altivelis* genome was carried out using the FALCON assembler^7^, followed by two rounds of polishing with Quiver^8^. FALCON implements a hierarchical assembly process that include the following steps: (1) subread error correction through aligning all reads to each other using daligner^9^, the overlap data were then processed to generate error-corrected consensus reads; after error correction, we obtained 28 Gb (35 X coverage) of error-corrected reads; (2) second round of overlap detection using error-corrected reads; (3) construction of a directed string graph from overlap data; and (4) resolving contig path from the string graph. After FALCON assembly, the genome was polished by Quiver. Initial assembly of the PacBio data resulted in a contig N50 (the minimum length of contigs accounting for half of the haploid genome size) of 4.14 Mb. Then, PacBio contigs were first scaffolded using optical map data, and the resulting scaffolds were further connected to super-scaffolds by 10X Genomics linked-read data using the fragScaff software^10^. Finally, we used Illumina-derived short reads to correct any remaining errors by pilon^11^. The final genome assembly of *C. altivelis* was with a total length of 1.07 Gb, contig N50 of 4,14 Mb, and scaffold N50 of 44.78 Mb (Table 2).

**Table 2.** Assembly statistics of *C. altivelis*.

| Sample ID | Length |  | Number |  |
| --- | --- | --- | --- | --- |
|  | Contig*(bp) | Scaffold(bp) | Contig* | Scaffold |
| Total | 1,063,656,417 | 1,066,981,559 | 1,256 | 534 |
| Max | 22,617,893 | 61,407,693 | - | - |
| Number≥2000 | - | - | 1,234 | 512 |
| N50 | 4,138,418 | 44,777,227 | 69 | 11 |
| N60 | 3,138,879 | 43,704,387 | 98 | 14 |
| N70 | 2,306,105 | 42,397,121 | 138 | 16 |
| N80 | 1,473,997 | 38,676,134 | 194 | 19 |
| N90 | 751,370 | 37,812,571 | 293 | 21 |

Note: The \* indicates contig after scaffolding.

Hi-C technology was further used for chromosome construction. We performed quality control of Hi-C raw data using HiCUP (version 3.0). We then aligned the raw reads to the draft assembled sequence by Bowtie2 (version 2.2.2), and filtered out the low quality reads to build raw intrachromosomal contact maps. Based on high quality Hi-C data, we anchored and orientated primary scaffolds into 24 chromosomes (Fig. 2), which additively covered 99.24% of the whole genome sequences.

**Figure 1.**
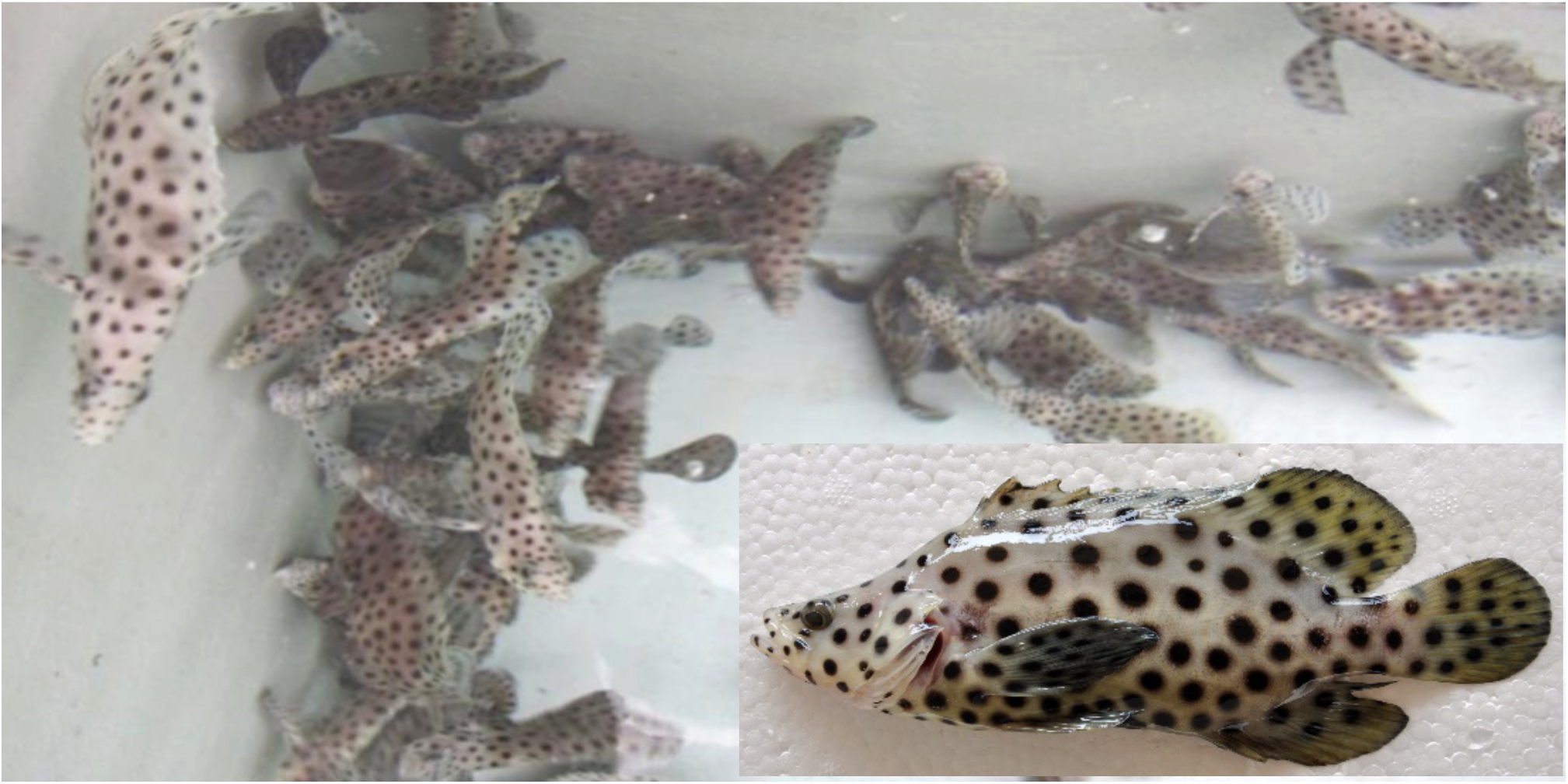
Morphology of the humpback grouper *C. altivelas*.

**Figure 2.**
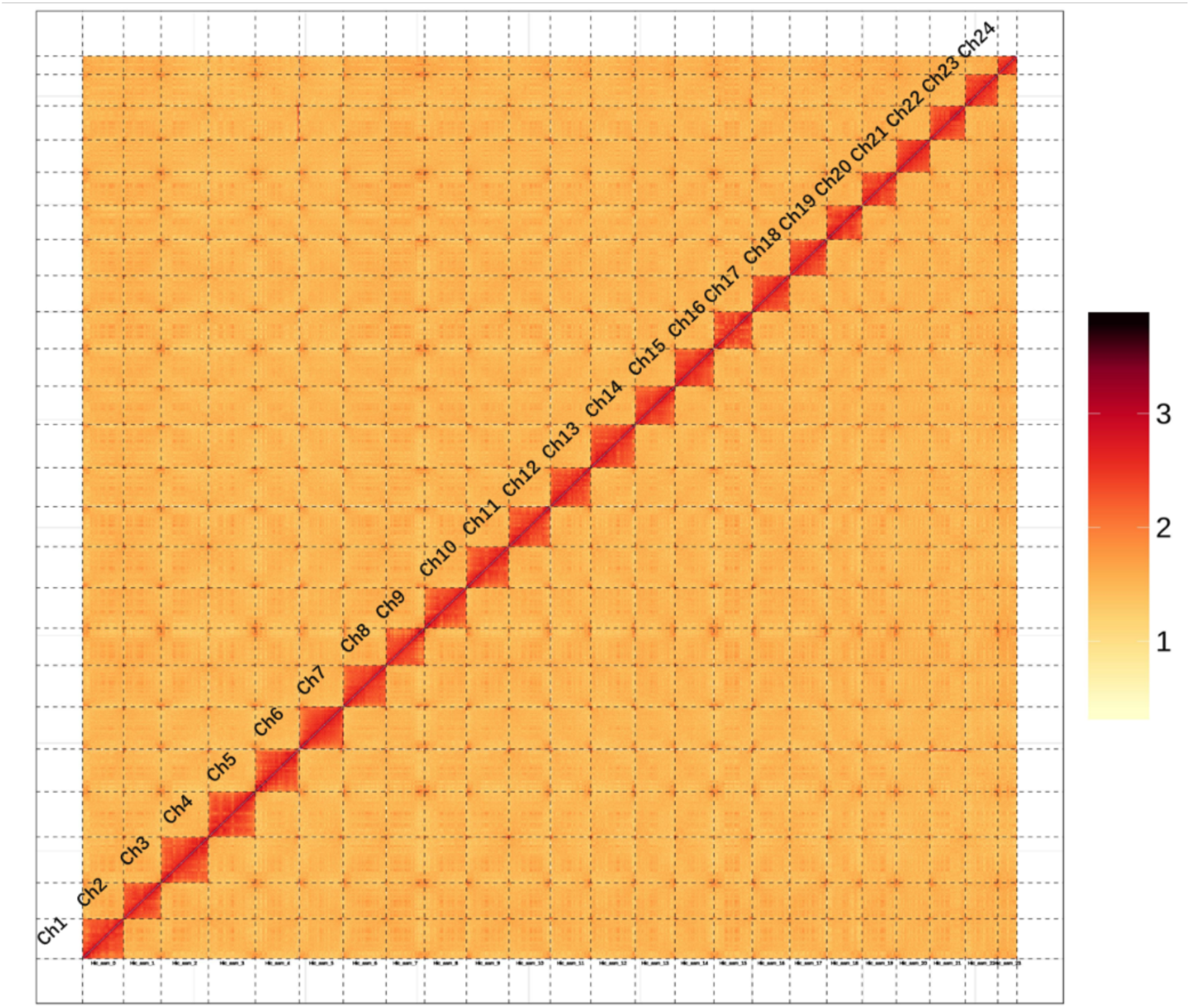
Hi-C chromosomal contact map of *C. altivelis*. The blocks represent the contacts between one location and another. The color reflects the intensity of each contact, with darker color indicates higher contact intensity.

### Repetitive sequences annotation

The repetitive elements in the *C. altivelis* genome were identified by a combination of evidence-based and *ab initio* approaches. We first used RepeatMasker (RepeatMasker, RRID:SCR 012954)^12^ and RepeatProteinMask to search against Repbase. We then construct a *de novo* repetitive element library using RepeatModeler and further utilized this *de novo* library for second round searching by RepeatMasker. In addition, we used Tandem Repeats Finder^13^, LTR FINDER (LTR FINDER, RRID:SCR 015247)^14^, PILER^15^, and RepeatScout (RepeatScout, RRID:SCR 014653)^16^ with default parameters for further repetitive elements annotation. Overall, we found 473,252,116 bp repeat sequences, accounted for 44.35% of *C. altivelis* genome (Table 3A), including 3.8% tandem repeats. Among transposable elements (TEs), there are 17.28% DNA transposons, 24.07% retroelements including LINE, SINE and LTR, and 3.74% unclassified elements (Table 3B).

**Table 3.**
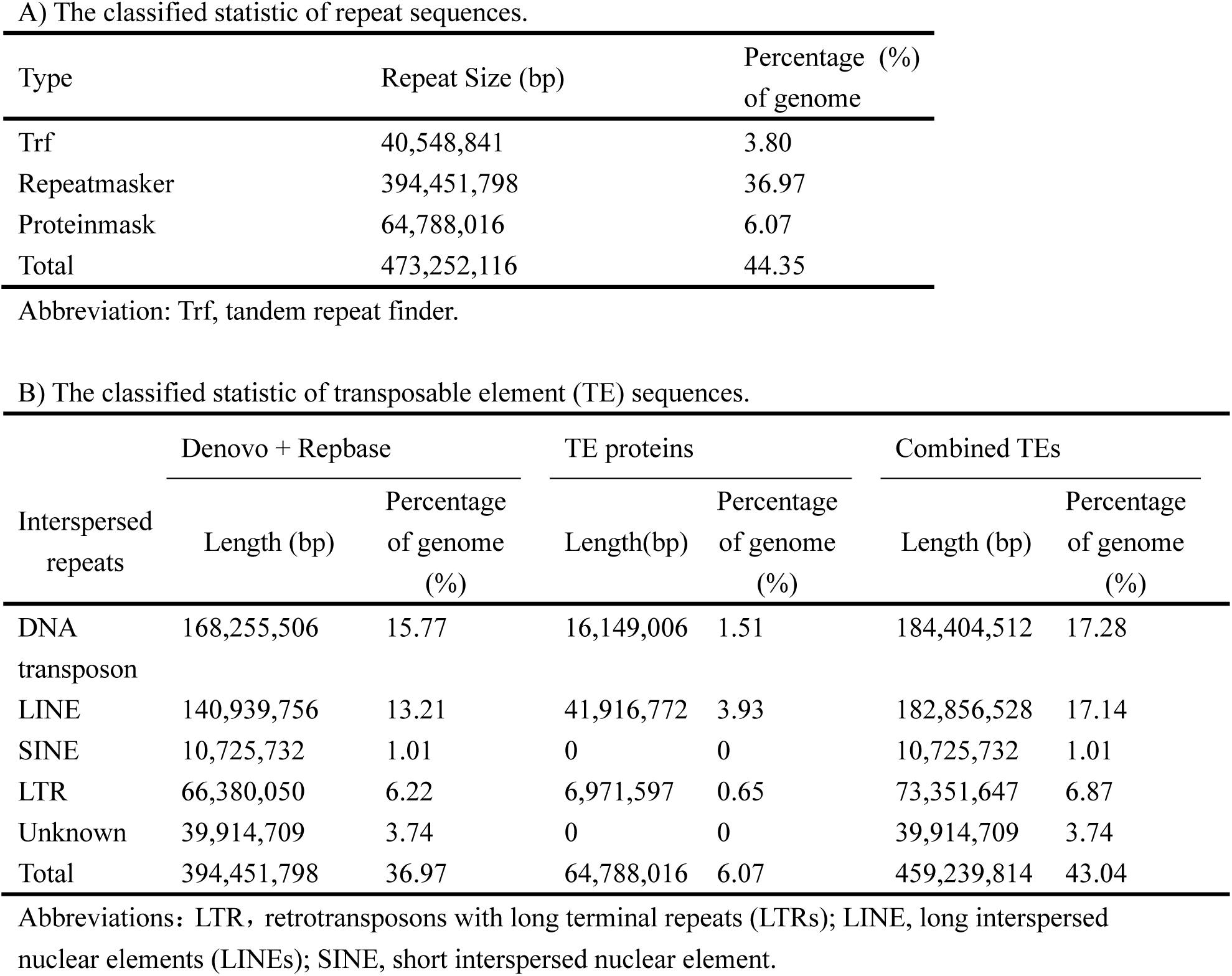
Summary of repetitive elements annotated in the genome of *C. altivelas*.

### Protein-coding gene prediction and functional annotation

To obtain a fully annotated *C. altivelas* genome, three approaches were combined to predict protein-coding genes including homology-based prediction, *ab initio* prediction, and transcriptome-based prediction. First, homology-based prediction was performed by TBLASTN (TBLASTN, RRID:SCR 011822)^17^ using protein repertoires of nine common vertebrates including *Branchiostoma floridae* (Bfl, GCA_000003815.1), *Cynoglossus semilaevis* (Cse, GCA_000523025.1), *Danio rerio* (Dre, GCF_000002035.6), *Gasterosteus aculeatus* (Gac, GCA_000180675.1), *Larimichthys crocea* (Lcr, GCA_000972845.1), *Oryzias latipes* (Ola, GCA_002234675.1), *Oreochromis niloticus* (Oni, GCF_001858045.1), and *Takifugu rubripes* (Tru, GCF_000180615.1). The Basic Local Alignment Search Tool (BLAST) hits were then conjoined by Solar software^18^. GeneWise (GeneWise, RRID:SCR 015054)^19^ was then used to predict the exact gene structure of the corresponding genomic region on each BLAST hit. Homology predictions were denoted as “Homology-set”.

Second, to provide further evidence for evaluating the predicted gene models, we assembled 38.67 Gb RNA-sequencing (RNA-seq) data derived from five different tissues by both *de novo* and reference-guided approaches. *De novo* RNA-seq assembly approach was performed by Trinity pipeline^20^, resulting in 370,688 contigs with an average length of 909 bp (Trinity-set). For reference-guided approach, short reads were directly mapped to the genome using Tophat (Tophat, RRID:SCR 013035)^21^ to identify putative exon regions and splice junctions. Cufflinks (Cufflinks, RRID:SCR 014597)^22^ and cuffmerge was then used to assemble the mapped reads into gene models (Cufflinks-set). These assembled Trinity-set and Cufflinks-set were then aligned against the *C. altivelis* genome by Program to Assemble Spliced Alignment (PASA). Valid transcript alignments were clustered based on genome mapping location and assembled into gene structures. Gene models created by PASA^23^ were denoted as “Transcripts-set”.

Third, *ab initio* prediction was performed on repeat-masked *C. altivelas* genome using Augustus (Augustus, RRID:SCR 008417)^24^, GeneID^25^, GeneScan^26^, GlimmerHMM (GlimmerHMM, RRID:SCR 002654)^27^ and SNAP^28^. Of these, Augustus, SNAP, and GlimmerHMM were trained by PASA-H-set gene models. Finally, three predicted gene models were integrated by EvidenceModeler^29^. Weights for each type of evidence were set as follows: Transdecoder > GeneWise = Cufflinks-set > Augustus > GeneID = SNAP = GlimmerHMM = GeneScan. The gene models were further updated by PASA2 to generate untranslated regions, alternative splicing variation information. Finally, a total of 27,242 protein-coding genes were obtained with a mean of 8.7 exons per gene (Table 4). The lengths of genes, coding sequence, introns, and exons in *C. altivelis* were comparable to those of closely related genomes (Supplementary Table S1).

**Table 4.** General statistics of predicted protein-coding genes.

| Gene set | Number | Average transcript length (bp) | Average CDS length (bp) | Average exons per gene | Average exon length (bp) | Average intron length (bp) |
| --- | --- | --- | --- | --- | --- | --- |
| *De novo* |  |  |  |  |  |  |
| Augustus | 34,508 | 12,616.88 | 1,249.14 | 6.90 | 181.02 | 1,926.57 |
| GlimmerHMM | 108,356 | 8,824.39 | 588.74 | 4.01 | 146.99 | 2,740.22 |
| SNAP | 45,011 | 31,206.98 | 1,053.25 | 7.54 | 139.75 | 4,613.14 |
| Geneid | 38,187 | 18,642.69 | 1,287.20 | 6.13 | 210.00 | 3,383.49 |
| Genscan | 39,221 | 19,619.96 | 1,492.24 | 8.05 | 185.36 | 2,571.11 |
| *Homolog* |  |  |  |  |  |  |
| Bfl | 20,910 | 5,887.97 | 814.08 | 4.14 | 196.55 | 1,614.99 |
| Cse | 30,239 | 11,878.94 | 1,448.76 | 6.76 | 214.25 | 1,810.18 |
| Dre | 31,724 | 9,717.79 | 1,252.27 | 5.92 | 211.60 | 1,721.32 |
| Gac | 38,491 | 8,349.92 | 1,023.10 | 5.28 | 193.89 | 1,713.23 |
| Lcr | 29,385 | 12,219.86 | 1,447.84 | 7.15 | 202.42 | 1,750.80 |
| Ola | 37,763 | 7,584.43 | 1,071.37 | 5.08 | 210.75 | 1,594.93 |
| Oni | 39,158 | 8,684.65 | 1,141.11 | 5.52 | 206.89 | 1,670.56 |
| Tru | 32,624 | 9,915.61 | 1,158.52 | 5.91 | 196.13 | 1,784.63 |
| *RNA-seq* |  |  |  |  |  |  |
| PASA | 74,899 | 16,509.17 | 1,322.94 | 8.02 | 165.03 | 2,164.37 |
| Cufflinks | 57,194 | 23,796.62 | 3,542.33 | 9.23 | 383.69 | 2,460.37 |
| *Integration* |  |  |  |  |  |  |
| EVM | 35,518 | 13,928.93 | 1,274.73 | 7.15 | 178.22 | 2,056.75 |
| Pasa-update* | 35,024 | 14,828.91 | 1,311.80 | 7.34 | 178.70 | 2,131.81 |
| Final set* | 27,242 | 17,687.21 | 1,510.89 | 8.71 | 173.37 | 2,096.83 |

Note: Items with \* stand for UTR region included, while other items did not include UTR regions.
Abbreviations: Bfl (*Branchiostoma floridae*), Cse (*Cynoglossus semilaevis*), Dre (*Danio rerio*), Gac (*Gasterosteus aculeatus*), Lcr (*Larimichthys crocea*), Ola (*Oryzias latipes*), Oni (*Oreochromis niloticus*), Tru (*Takifugu rubripes*).

Gene functions of protein-coding genes were annotated by searching functional motifs, domains, and the possible biological process of genes to known databases such as SwissProt^30^, Pfam ^31^, NR database (from NCBI), Gene Ontology^32^, and Kyoto Encyclopedia of Genes and Genomes^33^. A total of 27,067 protein-coding genes (99.4%) were successfully annotated for at least one function terms (Supplementary Table S2).

### Non-coding gene prediction

We also predicted noncoding RNA genes in the *C. altivelis* genome. The rRNA fragments were predicted by searching against human rRNA database using BLAST with an E-value of 1E-10. The tRNA genes were identified by tRNAscan-SE (tRNAscan-SE, RRID:SCR 010835) software^34^. The miRNA and snRNA genes were predicted by INFERNAL (INFERNAL, RRID:SCR 011809) ^35^ using Rfam database^36^. We found 410 ribosomal RNA (rRNA), 1,509 transfer RNA (tRNA), 1,335 microRNAs (miRNA), and 456 snRNA genes in the *C. altivelis* genome (Supplementary Table S3).

### Genome evolution analysis

To trace the evolutionary position of *C. altivelis*, nucleotide and protein datasets containing 1082 single-copy genes from the 16 species were used for phylogenetic tree reconstruction and divergence time estimation. The species included *Sillago sinica* (DOI:10.5524/100490), *Acanthopagrus schlegelii* (DOI:10.5524/100409), *L. crocea* (GCF_000972845.1), *O. latipes* (GCF_002234675.1), *O. niloticus* (GCF_001858045.1), *T. rubripes* (GCF_000180615.1), *D. rerio* (GCF_000002035.6), *Lepisosteus oculatus* (GCF_000242695.1), *Callorhinchus milii* (GCF_000165045.1), *Gasterosteus aculeatus* (GCA_000180675.1), *Gadus morhua* (GCA_000231765.1), *C. semilaevis* (GCF_000523025.1), *Xiphophorus maculate* (GCF_002775205.1), *Homo sapiens* (GCF_000001405.38), *Gallus gallus* (GCF_000002315.5) and *Ctenopharyngodon idella* (DOI: 10.5524/100494). All data were downloaded from NCBI or GigaDB database. To remove redundancy caused by alternative splicing variations, we retained only gene models at each gene locus that encoded the longest protein sequence. To exclude putative fragmented genes, genes encoding protein sequences shorter than 30 amino acids were filtered out.

Gene family analysis was conducted based on the homologs of protein-coding genes in the related species. All-against-all BLASTP (BLASTP, RRID:SCR 001010) was employed to identity the similarities among filtered protein sequences in these species with an E-value cutoff of 1e^-7^. The OrthoMCL (OrthoMCL, RRID:SCR 007839)^37^ method was used to cluster genes from these different species into gene families with the parameter of “-inflation 1.5”. Finally, a total of 23,140 gene family clusters were constructed. There were 1,045 gene families and 1,584 genes in *C. altivelis* without significant homologous hits to *L. crocea, L. oculatus* and *D. rerio*.

For phylogenetic analysis, MUSCLE (MUSCLE, RRID:SCR 011812)^38^ was used to generate multiple sequence alignments for protein sequences in each single-copy family with default parameters. Then, the alignments of each family were concatenated to a super alignment matrix. The super alignment matrix was used for phylogenetic tree reconstruction through maximum likelihood methods (Fig. 3). The clade with *H. sapiens and G. gallus* was set as outgroup.

**Figure 3.**
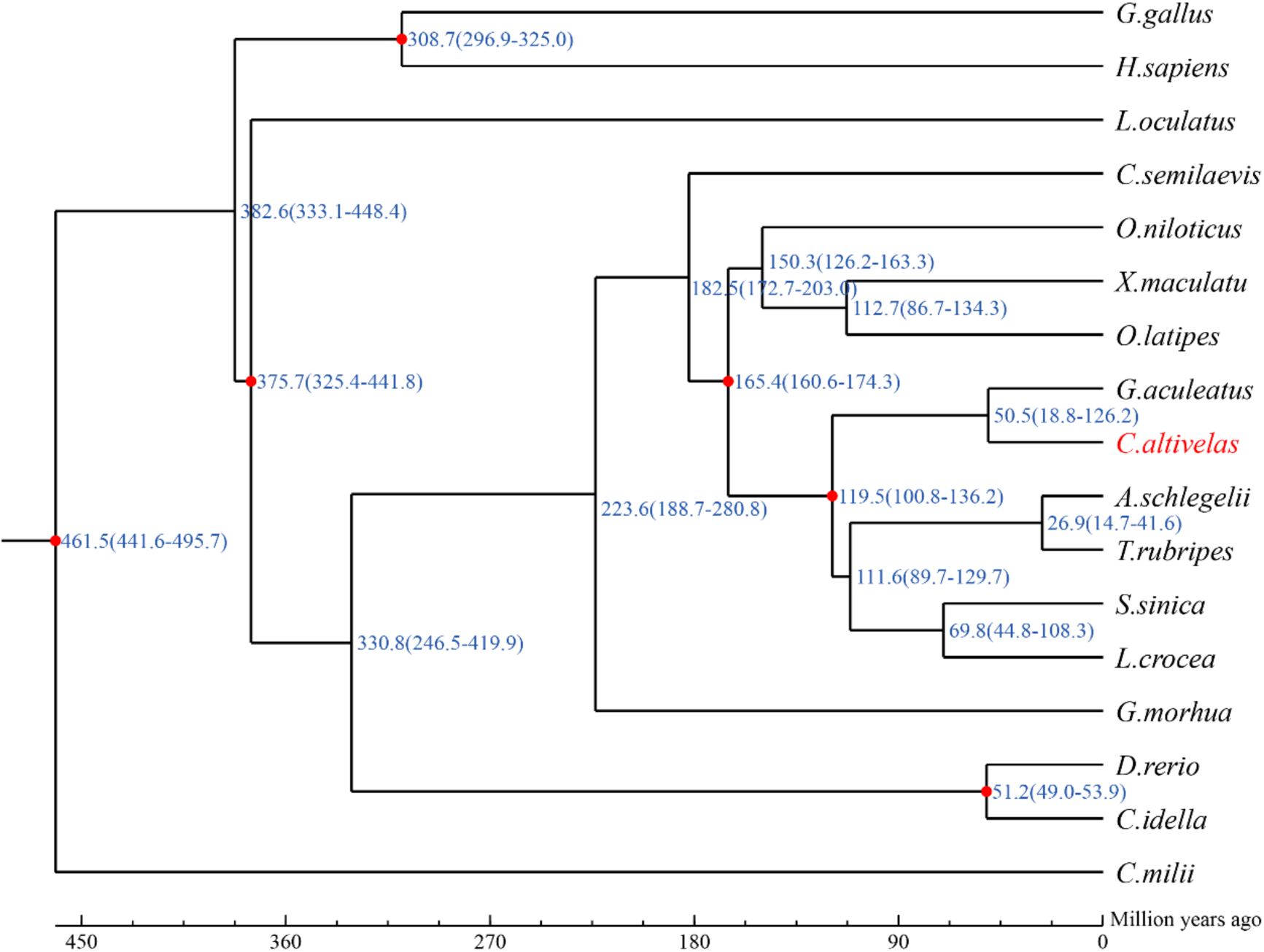
Divergence time estimated between *C. altivelis* and other species.

Divergence time was estimated based on the same dataset based on 1,082 single-copy genes from the 16 species using MCMCtree in PAML^39^ with the options “correlated molecular clock” and “JC69” model. A Markov chain Monte Carlo analysis was run for 20,000 generations using a burn-in of 1,000 iterations. Divergence time for the common ancestor of *C. milii* and *L. oculatus* (450∼497 Mya), *L. oculatus* and *C. idella* (291∼338 Mya), *T. rubripes* and *O. latipes* (163∼191 Mya), *G. aculeatus* and *T. rubripes* (101∼136 Mya), *C. idella* and *D. rerio* (49∼54 Mya), *H. sapiens and G. gallus* (292∼326 Mya) obtained from the TimeTree database (http://www.timetree.org/) and fossil records was used as the calibrate point. These phylogenetic analyses indicated that *C. altivelis* diverged from the common ancestral of *G. aculeatus* approximately 50.5 million years ago (Fig.3).

### Data Records

The sequenced raw data has been deposited in NCBI Sequence Read Archive (SRA) under BioProject accession PRJNA639378. The assembled chromosome level assembly, assembled contigs and annotation files are available in the figshare database (https://figshare.com/s/2d51c59fc548657f2ae8).

### Technical validation *of* the *C. altivelis* genome assembly

First, Illumina short reads were mapped to the *C. altivelis* genome with BWA^40^ (BWA, RRID: SCR 010910). The mapping rate is as high as 99.22% with a genome coverage of 99.64%. We further called and filtered single-nucleotide polymorphisms (SNPs) with SAMtools (SAMTools, RRID:SCR 002105)^41^. A total of 999,978 SNPs were identified including 997,151 heterozygous and 2,827 homozygous SNPs (Supplementary Table S4). The low rate of homozygous SNPs (0.0003% of the assembly) reflects a high-accuracy of genome assembly at the single base level.

Second, we assessed the completeness of the assembly with BUSCO^42^ and CEGMA^43^. Overall, 97.1% complete and 1.7% partial of the 2,586 vertebrate BUSCO genes were identified in the assembled genome. According to CEGMA, 226 (91.13%) complete matches and 235 (94.76%) complete plus partial matches of 248 core eukaryotic genes in CEGMA were identified in the genome assembly of *C. altivelis* genome.

## Code availability

No specific code was developed in this work. The data analyses were performed according to the manuals and protocols provided by the developers of the corresponding bioinformatics tools that are described in the Methods section.

## Acknowledgements

This research was supported financially by the National Natural Science Foundation of China (No. 41666006), Natural Science Foundation of Hainan Province (No. 2019RC078), and the Nanhai Famous Youth Project.

## Author contributions

Y.S. and Y.Z. designed research; Y.S., D.Z., J.S., G.C., Y.W., Y.S., Z.C., and L.Z. analyzed data; Y.S, L.Z., D.Z. J.S., and Y.Z. wrote the manuscript; and all authors read, edited, and approved the final manuscript.

## Abbreviations

BLAST: Basic Local Alignment Search Tool
BUSCO: Benchmarking Universal Single-Copy Orthologs
CEGMA: Core Eukaryotic Genes Mapping Approach
MITE: miniature inverted–repeat transposable elements
NCBI: National Center for Biotechnology Information
PacBio: Pacific Biosciences
PASA: Program to Assemble Spliced Alignment
PASA-T-set: PASA Trinity set
RNA-seq: RNA sequencing

## Additional files

Supplemental file.pdf

## References

1. M. W. Johnston, S. J. Purkis, Modeling the potential spread of the recently identified non-native panther grouper (*Chromileptes altivelis*) in the Atlantic using a cellular automaton approach. PloS one 8, 1–11 (2013).

2. K. Williams, D. Smith, I. Williams, S. Irvin, M. Barclay, ACIAR grouper grow-out feeds program and related CSIRO research. Aquaculture Asia magazine 10, 29–33 (2005).

3. J. Qin, D. Hu, W. Yang, J. Xiao, Complete mitochondrial genome of the humpback grouper *Cromileptes altivelis*. Mitochondrial DNA 25, 200–201 (2014).

4. K. Mahardika, A. Yamamoto, T. Miyazaki, Susceptibility of juvenile humpback grouper *Cromileptes altivelis* to grouper sleepy disease iridovirus (GSDIV). Diseases of aquatic organisms 59, 1–9 (2004).

5. K. Williams, S. Irvin, M. Barclay, Polka dot grouper *Cromileptes altivelis* fingerlings require high protein and moderate lipid diets for optimal growth and nutrient retention. Aquaculture nutrition 10, 125–134 (2004).

6. G. Marçais, C. Kingsford, A fast, lock-free approach for efficient parallel counting of occurrences of k-mers. Bioinformatics 27, 764–770 (2011).

7. M. Pendleton et al., Assembly and diploid architecture of an individual human genome via single-molecule technologies. Nature methods 12, 780 (2015).

8. C.-S. Chin et al., Nonhybrid, finished microbial genome assemblies from long-read SMRT sequencing data. Nature methods 10, 563 (2013).

9. G. Myers, in International Workshop on Algorithms in Bioinformatics. (Springer, 2014), pp. 52–67.

10. A. Adey et al., In vitro, long-range sequence information for de novo genome assembly via transposase contiguity. Genome research 24, 2041–2049 (2014).

11. G. Parra, K. Bradnam, I. Korf, CEGMA: a pipeline to accurately annotate core genes in eukaryotic genomes. Bioinformatics 23, 1061–1067 (2007).

12. C. M. Bergman, H. Quesneville, Discovering and detecting transposable elements in genome sequences. Briefings in bioinformatics 8, 382–392 (2007).

13. G. Benson, Tandem repeats finder: a program to analyze DNA sequences. Nucleic acids research 27, 573–580 (1999).

14. Z. Xu, H. Wang, LTR_FINDER: an efficient tool for the prediction of full-length LTR retrotransposons. Nucleic acids research 35, 265–268 (2007).

15. R. C. Edgar, E. W. Myers, PILER: identification and classification of genomic repeats. Bioinformatics 21, 152–158 (2005).

16. A. L. Price, N. C. Jones, P. A. Pevzner, De novo identification of repeat families in large genomes. Bioinformatics 21, 351–358 (2005).

17. S. F. Altschul, W. Gish, W. Miller, E. W. Myers, D. J. Lipman, Basic local alignment search tool. Journal of molecular biology 215, 403–410 (1990).

18. X.-J. Yu, H.-K. Zheng, J. Wang, W. Wang, B. Su, Detecting lineage-specific adaptive evolution of brain-expressed genes in human using rhesus macaque as outgroup. Genomics 88, 745–751 (2006).

19. E. Birney, M. Clamp, R. Durbin, GeneWise and genomewise. Genome research 14, 988–995 (2004).

20. M. G. Grabherr et al., Full-length transcriptome assembly from RNA-Seq data without a reference genome. Nature biotechnology 29, 644 (2011).

21. D. Kim et al., TopHat2: accurate alignment of transcriptomes in the presence of insertions, deletions and gene fusions. Genome biology 14, R36 (2013).

22. C. Trapnell et al., Differential gene and transcript expression analysis of RNA-seq experiments with TopHat and Cufflinks. Nature protocols 7, 562–578 (2012).

23. B. J. Haas et al., Improving the Arabidopsis genome annotation using maximal transcript alignment assemblies. Nucleic acids research 31, 5654–5666 (2003).

24. M. Stanke, S. Waack, Gene prediction with a hidden Markov model and a new intron submodel. Bioinformatics 19, 215–225 (2003).

25. R. Guigo, Assembling genes from predicted exons in linear time with dynamic programming. Journal of Computational Biology 5, 681–702 (1998).

26. C. Burge, S. Karlin, Prediction of complete gene structures in human genomic DNA. Journal of molecular biology 268, 78–94 (1997).

27. W. H. Majoros, M. Pertea, S. L. Salzberg, TigrScan and GlimmerHMM: two open source ab initio eukaryotic gene-finders. Bioinformatics 20, 2878–2879 (2004).

28. I. Korf, Gene finding in novel genomes. BMC bioinformatics 5, 59 (2004).

29. B. J. Haas et al., Automated eukaryotic gene structure annotation using EVidenceModeler and the Program to Assemble Spliced Alignments. Genome biology 9, 1–22 (2008).

30. R. Apweiler et al., UniProt: the universal protein knowledgebase. Nucleic acids research 32, 115–119 (2004).

31. R. D. Finn et al., The Pfam protein families database: towards a more sustainable future. Nucleic acids research 44, 279–285 (2016).

32. G. O. Consortium, Expansion of the Gene Ontology knowledgebase and resources. Nucleic acids research 45, 331–338 (2017).

33. T. M. Lowe, S. R. Eddy, tRNAscan-SE: a program for improved detection of transfer RNA genes in genomic sequence. Nucleic acids research 25, 955–964 (1997).

34. E. P. Nawrocki, D. L. Kolbe, S. R. Eddy, Infernal 1.0: inference of RNA alignments. Bioinformatics 25, 1335–1337 (2009).

35. Y.-h. Li et al., De novo assembly of soybean wild relatives for pan-genome analysis of diversity and agronomic traits. Nature biotechnology 32, 1045 (2014).

36. M. Kanehisa et al., Data, information, knowledge and principle: back to metabolism in KEGG. Nucleic acids research 42, 199–205 (2014).

37. L. Li, C. J. Stoeckert, D. S. Roos, OrthoMCL: identification of ortholog groups for eukaryotic genomes. Genome research 13, 2178–2189 (2003).

38. R. C. Edgar, MUSCLE: multiple sequence alignment with high accuracy and high throughput. Nucleic acids research 32, 1792–1797 (2004).

39. Z. Yang, PAML: a program package for phylogenetic analysis by maximum likelihood. Bioinformatics 13, 555–556 (1997).

40. B. J. Walker et al., Pilon: an integrated tool for comprehensive microbial variant detection and genome assembly improvement. PloS one 9, e112963 (2014).

41. H. Li, R. Durbin, Fast and accurate short read alignment with Burrows–Wheeler transform. bioinformatics 25, 1754–1760 (2009).

42. H. Li, A statistical framework for SNP calling, mutation discovery, association mapping and population genetical parameter estimation from sequencing data. Bioinformatics 27, 2987–2993 (2011).

43. F. A. Simão, R. M. Waterhouse, P. Ioannidis, E. V. Kriventseva, E. M. Zdobnov, BUSCO: assessing genome assembly and annotation completeness with single-copy orthologs. Bioinformatics 31, 3210–3212 (2015).

