## Supplemental file for "A chromosome-level genome assembly and annotation of the humpback grouper *Cromileptes altivelas*"

Figure S1

The *k*-mer analysis (k=17) for estimating genome size of *C. altivelas*. The x-axis refers to the *k*-mer depth; the y-axis refers to the frequency of *k*-mer for a given depth.

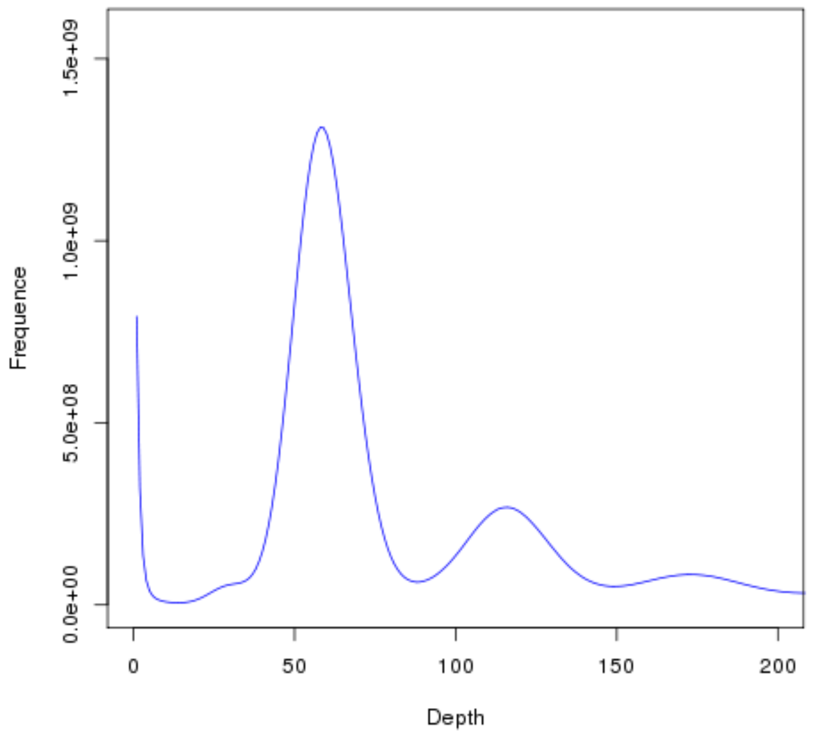

Table S1
The lengths of genes, coding sequence, introns, and exons in *C. altivelis* were comparable to those of closely related genomes

| Species | Number | Average transcript length(bp) | Average CDS length(bp) | Average exons per gene | Average exon length(bp) | Average intron length(bp) |
|---|---|---|---|---|---|---|
| Bfl | 28,621 | 9,236.65 | 1,392.24 | 7.03 | 198.11 | 1,301.41 |
| Cse | 21,256 | 11,471.84 | 1,791.91 | 10.67 | 168 | 1,001.42 |
| Dre | 25,619 | 25,207.59 | 1,642.64 | 9.42 | 174.39 | 2,798.97 |
| Gac | 20,787 | 8,451.06 | 1,548.67 | 10.4 | 148.94 | 734.44 |
| Lcr | 28,544 | 14,076.04 | 1,807.56 | 11.15 | 162.18 | 1,209.25 |
| Ola | 19,699 | 12,145.58 | 1,515.82 | 10.25 | 147.82 | 1,148.61 |
| Oni | 21,437 | 14,903.11 | 1,714.22 | 10.9 | 157.25 | 1,332.07 |
| Tru | 18,523 | 7,492.75 | 1,693.53 | 11.1 | 152.61 | 574.33 |

Abbreviations: Bfl (*Branchiostoma floridae*), Cse (*Cynoglossus semilaevis*), Dre (*Danio rerio*), Gac (*Gasterosteus aculeatus*), Lcr (*Larimichthys crocea*), Ola (*Oryzias latipes*), Oni (*Oreochromis niloticus*), Tru (*Takifugu rubripes*).

Table S2 Functional annotation for protein-coding genes in the genome of *C. altivelis* genome.

| Database | Number | Percentage (%) |
|---|---|---|
| Swissprot | 23,280 | 85.5 |
| Nr | 25,724 | 94.4 |
| KEGG | 21,902 | 80.4 |
| InterPro | 26,990 | 99.1 |
| GO | 25,117 | 92.2 |
| Pfam | 21,427 | 78.7 |
| Annotated | 27,067 | 99.4 |
| Unannotated | 175 | 0.6 |
| Total | 27,242 | - |

Table S3 Detailed results of ncRNA annotation.

| Type | | Copy (w*) | Average length (bp) | Total length (bp) | Percentage (%) of genome |
|---|---|---|---|---|---|
| miRNA | | 1,335 | 127.24 | 169,865 | 0.01592 |
| tRNA | | 1,509 | 76.01 | 114,705 | 0.01075 |
| rRNA | rRNA | 410 | 149.49 | 61,292 | 0.005744 |
| | 18S | 109 | 154.39 | 16,828 | 0.001577 |
| | 28S | 189 | 172.34 | 32,573 | 0.003053 |
| | 5.8S | 3 | 125.67 | 377 | 0.000035 |
| | 5S | 109 | 105.63 | 11,514 | 0.001079 |
| snRNA | snRNA | 456 | 153.17 | 69,845 | 0.006546 |
| | CD-box | 219 | 145.33 | 31,827 | 0.002983 |
| | HACA-box | 94 | 157.59 | 14,813 | 0.001388 |
| | Spliceosomal RNA | 75 | 146.79 | 11,009 | 0.001032 |

Note: The w* stands for whole genome.

Table S4 The summary of SNPs of *C. altivelas* genome assembly.

| Type | Number | Percentage (%) of genome |
|---|---|---|
| All SNP | 999978 | 0.09 |
| Heterozygosis SNP | 997151 | 0.09 |
| Homology SNP | 2827 | 0.0003 |
